# Path vectors: a neural code for sequential memory

**DOI:** 10.1101/2022.02.28.482342

**Authors:** Rich Pang, Stefano Recanatesi

## Abstract

While recalling lists of unrelated items is highly challenging we can recall much longer sequences structured as an episode or story. It is unknown why these patterns have such a striking influence on memory. We introduce a model where the experience of an episode or story is represented as a path through a pre-existing network of cognitive states. We demonstrate that by summing the neural representations of the visited states, this path can be transformed into a simple neural code: a path vector. We show how, by leveraging sparse connectivity and high dimensionality, path vectors provide robust codes for a large set of sequences and can be decoded mechanistically for memory retrieval. Fitting our model to data reveals how human free and serial recall may emerge from adapting coding mechanisms tuned for sequences aligned with existing network paths. We thus posit that sequences such as episodes or stories map more directly to existing cognitive network paths than arbitrary lists, with the latter eliciting paths that tend to interfere and impair recall. Our model suggests that mnemonic strategies like imposing narrative structure on a list act to improve recall by reducing such path interference. This work illuminates a simple bioplausible means for flexibly recruiting existing cognitive structures to encode new memories.

---

Episodic memory, the storage and recall of specific past experiences, is indispensable to cognition and behavior. Typically, an episode contains a *novel sequence* of events, like the sequence of actions taken to visit a new restaurant. The neural basis of remembering sequential episodes, especially those occurring only once, is unknown. A challenge is that sequences are not only the sum of their parts but also contain the ordering of the events or items. Although reflections of past event orderings in neural activity have long been studied (1–3), we still lack a deep understanding of how ordering information is distributed across neurons at any moment in time through which the memory persists.

Although a comprehensive picture has not emerged, models for sequence memory abound at many levels. Biophysically, sequences could be stored in synaptic changes (4–6) or graded intracellular states reactivated by oscillations (1, 7). From the viewpoint of distributed coding, models such as the temporal context model or vector symbolic architectures (8, 9) encode sequences by binding high-dimensional “item vectors” to positional or context vectors via e.g. multiplicative or convolution operations. Recurrent neural networks can also be trained to store input sequences via persistent activity (10), which can similarly lead to item-context vector representations of ordering, although trained networks are typically challenging to fully understand (11). Finally, high-level models, such as positional or chaining models (12–16) replicate certain human recall patterns but usually lack an underlying neural code. Beyond uncertainties over the cellular mechanisms these models require (e.g. synaptic changes for storing sequences may be too weak (17, 18) for one-time episodes, and oscillation models usually require precision timing (1, 19), challenging their robustness), none yet systematically addresses the remarkable variability and flexibility of memory. For instance, we recall stories better than random lists (20–23) and can even endow random lists with auxiliary narrative structure on the fly to improve our recall (24). Whether such complex yet deeply relatable phenomena can be described by a transparent mathematical model based on bioplausible neural codes is not clear.

Here we introduce such a model. We first convert a sequence of events or items to a path through a pre-existing network of cognitive states, then write this path via a simple sum into a neural code called a *path vector*. This yields a distributed, fixed-dimension representation from which the path and original sequence can later be retrieved. We find our model fits human free-recall data (where ordering need not be retained) but can also explain how people exploit existing cognitive paths to store novel ordering. By considering a network whose links reflect the transitional structure of everyday cognitive experiences, we posit that “natural” sequences, such as episodes or stories, better align with existing network paths and have fewer collisions than arbitrary sequences, improving recall. Although we do not attempt a comprehensive account of human memory, which may rely on many mechanisms, our model provides an interpretable hypothesis for flexible sequential memory. Moreover, it suggests a neurocognitive reason episodes and stories are more memorable and can be employed for mnemonic enhancement. This work may help us understand how the brain processes the sequential structures of everyday life and inform theories and treatments of memory disorders.

## Significance

How does the brain store memories of everyday experiences? And why are these so memorable compared to random lists? To address these questions we developed a theoretical framework based on networks of cognitive states and distributed brain activity. In our framework, experiences map to cognitive paths and are stored in memory as the sum of the neural patterns they activate–a path vector. Recall accuracy is shaped by how well experiences align with natural cognitive paths. We show how this not only explains core free and serial recall phenomena but also provides a simple, bio-plausible account of the vast flexibility of memory–including how auxiliary storylines can improve recall–based on creating and retracing paths through structured mental spaces.

## Results

### Path vector representation of typical event sequences

Many event sequences can be stored as the *set* of events alone, with their order retrievable from prior knowledge of how event sequences usually unfold. If morning outings usually consist of grabbing keys, choosing transportation, going somewhere, and performing a task; there is only one order in which {grab keys, ride bike, visit deli, buy coffee} could have occurred (Fig. 1a), so the brain need not additionally store the ordering. If the mental or *cognitive states* we transition through in daily life are similarly constrained, a sequence of such states can be stored as their set alone as well, with their order recoverable from prior knowledge of the typical transitions. Here we model such constraints via a *cognitive network* (Fig. 1b), whose nodes {*v*^*i*^} are cognitive states and whose connections *W*_*ij*_ are typical state transitions, or “associations”. If cognitive states and their associations correspond to events and their typical transitions (Figs. 1a to 1b) – as if the brain has previously learned a model of the world (25) – many everyday event sequences could be stored simply as the set of cognitive states 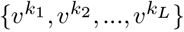 they evoke and serially retrieved by retracing the path 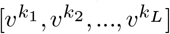 through the cognitive network that connects them (if the path contains no cycles.)

**Fig.1.**
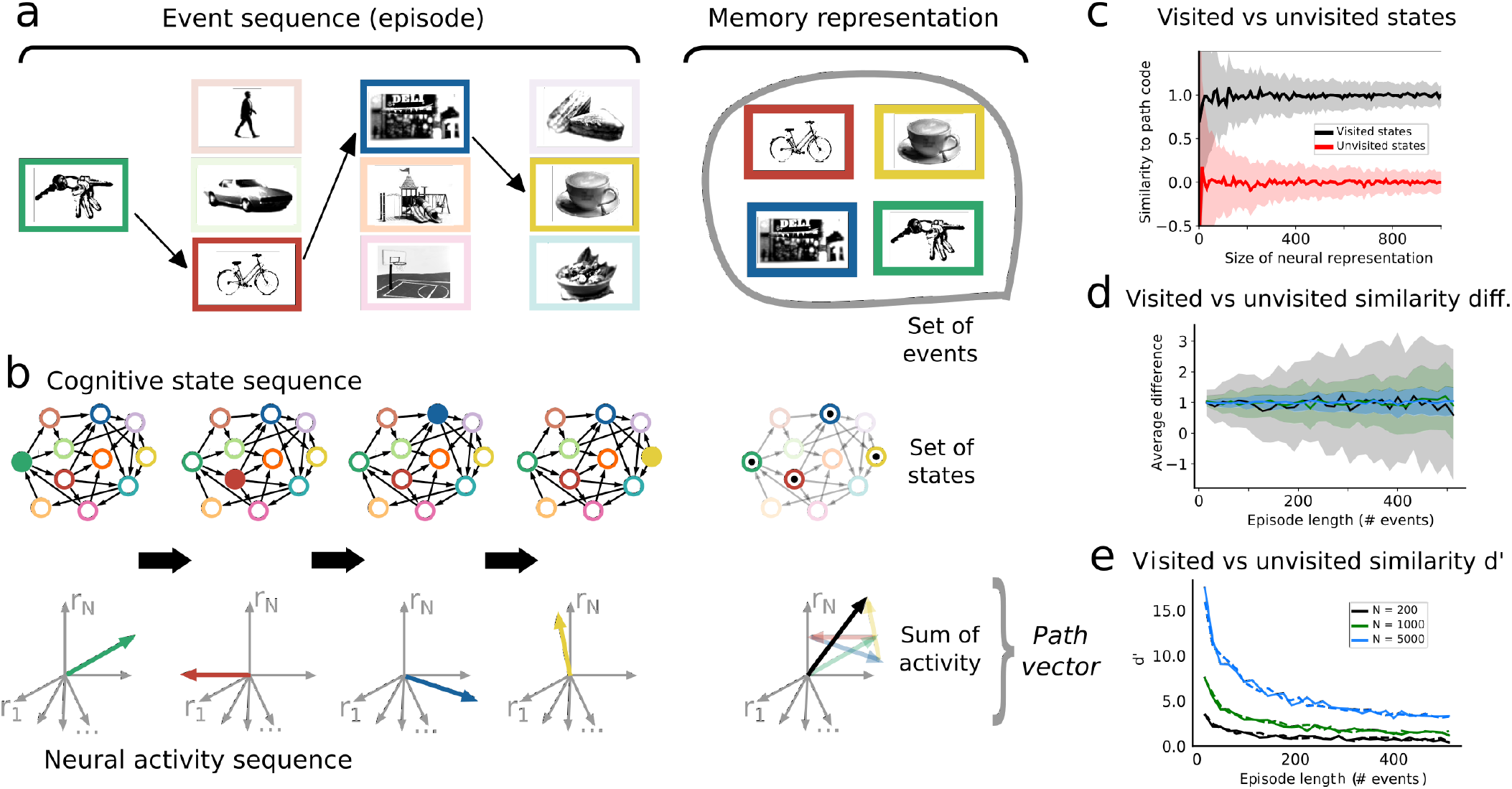
Path vector representation of a sequential experience. A. Left: Event sequence corresponding to a morning outing. Columns show possible events at different times; highlights show chosen sequence. Right: memory representation of the event sequence, i.e. just the set of events. B. Top left: a network of cognitive states and their associations, corresponding to the events in A. Filled circles indicate active state at each time. Top right: memory representation of cognitive state sequence, i.e. the set of states visited (inscribed dots). Bottom left: high-dimensional neural representation of cognitive states, with each axis depicting one neuron’s firing rate. Bottom right: stored memory of the neural activity, i.e. the sum of activated neural representations, called a *path vector*. C. Dot-product similarity of path vector to cognitive states either visited or unvisited during the event sequence (*L* = 16 events), vs number of neurons (computed over 100 instantiations of a network of *M* = 1000 states). Thick: mean; shading: STD. D. Difference between visited and unvisited states’ dot-product similarity to the path vector vs number of events/states visited in the sequence. Thick: mean; shading: STD. E. d’ between distributions of dot-product similarities with the path vector for either visited or unvisited states vs number of events/states visited. Computed over 100 instantiations of a network with either 1000 (solid) or 100,000 cognitive states (dashed).

We model the neural representation of each cognitive state *v*_*i*_ as a random population vector **r**_*i*_. This resembles how hippocampal populations with random place fields encode locations (26) or how olfactory circuits encode odors (27). For now we assume components of **r**_*i*_ are i.i.d., and such that E[‖**r**_*i*_‖] = 1. When the dimension (number of neurons) *N* is large, all state vectors will be nearly orthogonal (for exponential states in *N*) (9). Thus, out of *M* total states a subset of *L* of them visited in the experience be stored as the sum, or *path vector* (Fig. 1b),

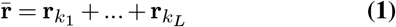

since 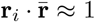 if 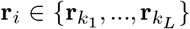 and 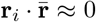 if not, with errors scaling as 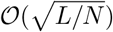. Visited vs unvisited states are more distinguishable when *L* ≪ *N*, but even a few thousand neurons accurately stores hundreds of states (Figs. 1c to 1e). Thus 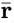 encodes the states, which in turn encode the path. A sequential episode aligning with a unique path through the cognitive state network can therefore be stored as a fixed-dimension path vector 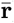, with all information evenly distributed across all neurons.

### Distinct regimes for free and natural recall

We model retrieval of the sequence from the path vector probabilistically. We call the initial experience of the sequence the *presentation phase*; events (or “items”, as in memory experiments) are shown serially to the model, activating a sequence of cognitive states, with neural representations 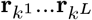, that are summed to form 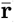. In the *retrieval phase*, cognitive states are traversed via

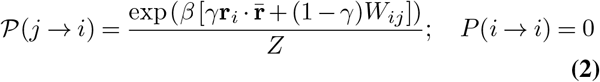

where *i* and *j* index the current and previous cognitive state. We say item *i* is retrieved when state *i* is visited. *W*_*ij*_ is binary, indicating whether a link from *j* → *i* exists in the network; *Z* normalizes *P* (*j* → *i*) such that ∑_*i*_*P*(*j* → *i*) = 1 *β* ∈ [0, ∞) controls retrieval stochasticity; and *γ* controls how strongly the network (*γ* = 0) vs the items “primed” by the path vector (*γ* = 1) influence retrieval (Fig. 2b).

**Fig.2.**
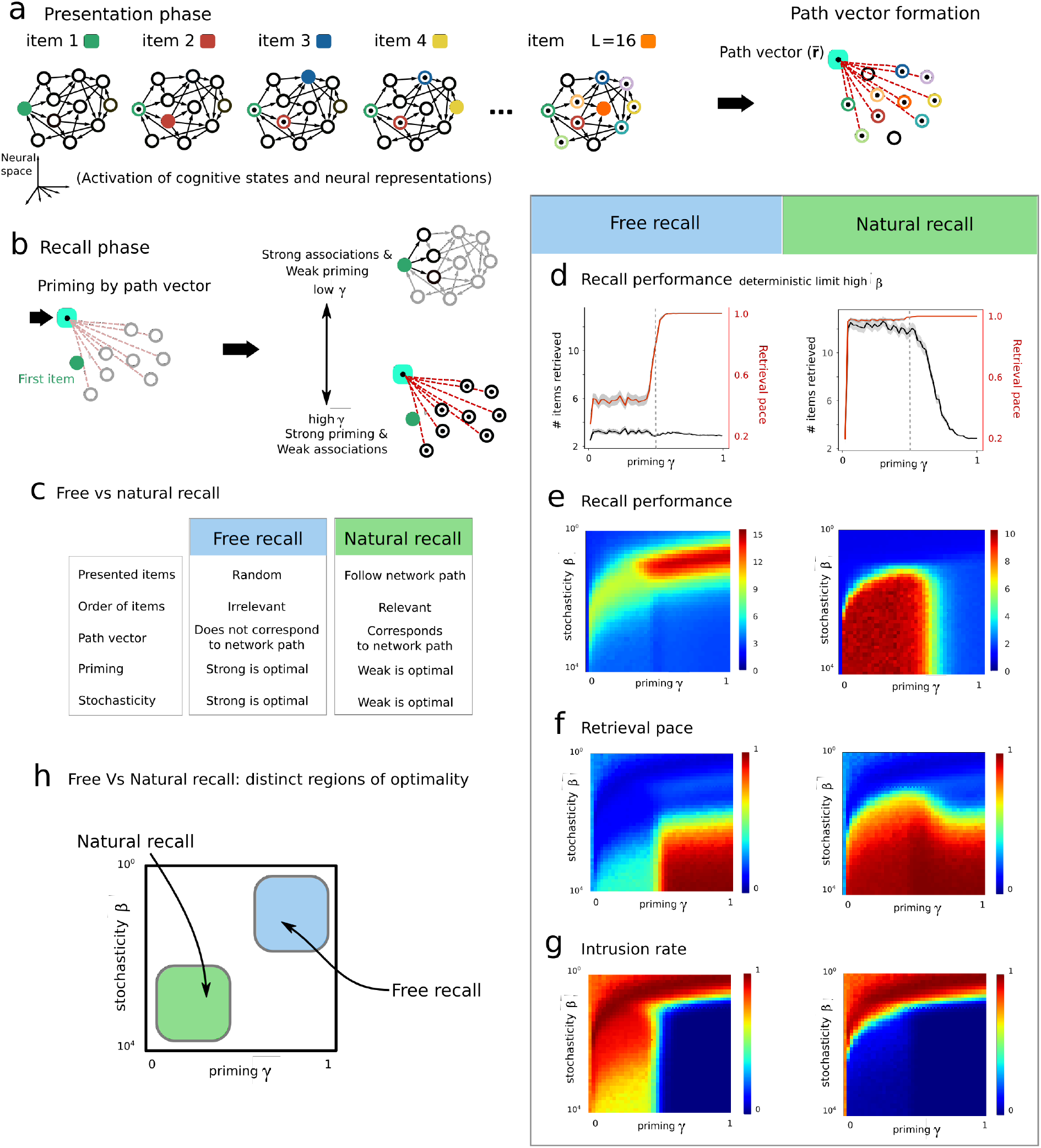
Recall of natural vs random sequences. a) Presentation phase of a sequence of items, where different items have different colors. During presentation the corresponding cognitive state (full color) and neural representation are activated. By the end of the presentation a path vector (cyan) is formed encoding for the presented items. b) The recall phase of the presented list is instantiated by the “reactivation” of the path vector. Successive items are recalled according to Equation 2, whose dynamics are regulated by *γ* (priming) and *β* (stochasticity). c) Comparison of free vs natural recall. d) Our model’s recall performance and retrieval pace vs priming *γ* in free and natural recall paradigms in the deterministic limit (*β* → ∞). e) Recall performance vs *γ, β*. f) Retrieval pace vs *γ, β*. g) Intrusion rate vs *γ, β*. h) Approximate region of optimality for free and natural recall.

We test this model on two memory tasks. In *free recall*, a standard task, items are presented randomly and the model must recall as many as possible in any order. In *natural recall*, a task designed for our model, (1) the presented item order follows the network *W*, (i.e. *W*_*ij*_ = 1 between consecutive items); (2) retrieval begins at the correct first item (we address later how to autonomously recall the first item); and (3) the sequence must be retrieved in the correct order. Natural recall generalizes Fig. 1’s example and resembles the experimental paradigm of serial recall (28), except that here item presentation follows network paths (Fig. 2c).

In each task we assess the roles of stochasticity (*β*) and priming (*γ*). Using a cognitive network of M=1000 states, corresponding to 1000 unique items, (*W*_*ij*_ = 1 with probability 0.01), we represent each state *v*_*i*_ by a random N=500 dimensional neural vector **r**_*i*_. In each trial we present L=16 out of the possible items and stop retrieval when 100 states/items are recalled (including repeats). We measure performance (number of correct items retrieved, with correct order required in natural recall), retrieval pace (total unique items retrieved divided by the timestep the last unique item was retrieved), and intrusion rate (number of incorrect over total items retrieved).

In the deterministic limit (*β* → ∞ ; Fig. 2d) a given item *j* is always followed by the same item *i*. In this limit natural recall outperforms free recall. When *γ* = 0 (no priming by the path vector) recall simply follows the underlying graph *W*, regardless of which items were presented, which is suboptimal in both tasks. As *γ* increases, the path vector more strongly primes the presented sequence, enabling close to perfect retrieval in natural recall. When priming grows too strong (*γ* → 1), however, recall dynamics gets trapped in the states most similar to the path vector, independent of *W*, so few states are visited and items retrieved. Thus, low stochasticity and moderate priming by the path vector strongly support natural but not free recall.

Varying both *β* and *γ* revealed disjoint regimes of optimal performance for each task (Figs. 2e to 2h). Free recall was optimized at low *β* (high noise) and *γ >* 0.5 while natural recall was optimized at high *β* (low noise) and *γ* < 0.5 but nonzero (Figs. 2e to 2g). Although good performance in free recall can emerge by strongly priming all presented words then randomly sampling among them (high *γ* low *β*, Fig. 2e), randomly hopping among items is slow (many already-retrieved items will be revisited), requiring a large number of timesteps to retrieve all items (Fig. 2f). In free recall performance is optimized at high noise and strong priming, but speed is optimized by exploiting the cognitive network (high *β*), highlighting a speed-performance trade-off absent in natural recall. For free recall, low intrusion rates required high priming, whereas the same in natural recall only required low noise (Fig. 2g), further supporting disjoint optimal parameter regimes for each task (Fig. 2h).

### Modeling human memory experiments

Which model parameters best reproduce human recall? As experiments cannot easily replicate our natural recall task, which requires knowing subjects’ hidden cognitive networks, we analyzed free recall experiments (29). In each trial we examined (141 human subjects, 112 trials each) a subject was visually shown 16 random words (Methods) (29). Since the true cognitive associations are unknown, we modeled *W* using pre-computed Latent Semantic Analysis (LSA) scores between pairs of presented words (Methods); LSA scores are higher for semantically related and lower for unrelated words. To assess which model parameters better reflected human free recall we varied *β* and *γ* and compared features of the model behavior to those in the data.

Human free-recall statistics were best reproduced when our model’s parameters were also optimized for free recall. The distribution of subjects’ performance (measured as how many correct words were recalled irrespective of order) was most similar to the data when the model operated with low *β* and high *γ* (top-right gray box, Fig. 3a); as was the case for the distribution of intrusion rates (Fig. 3b). The average LSA score between consecutively retrieved words (Fig. 3c) was also closest to the data in the same parameter region, suggesting that even in the free-recall parameter region, model retrieval dynamics are impacted by an underlying cognitive network, reflecting the known influence of semantic similarity in human free recall (30). This validates the model’s capacity to capture the free recall task, suggesting subjects might employ path-vector-like retrieval to recall items even when order is not required.

**Fig.3.**
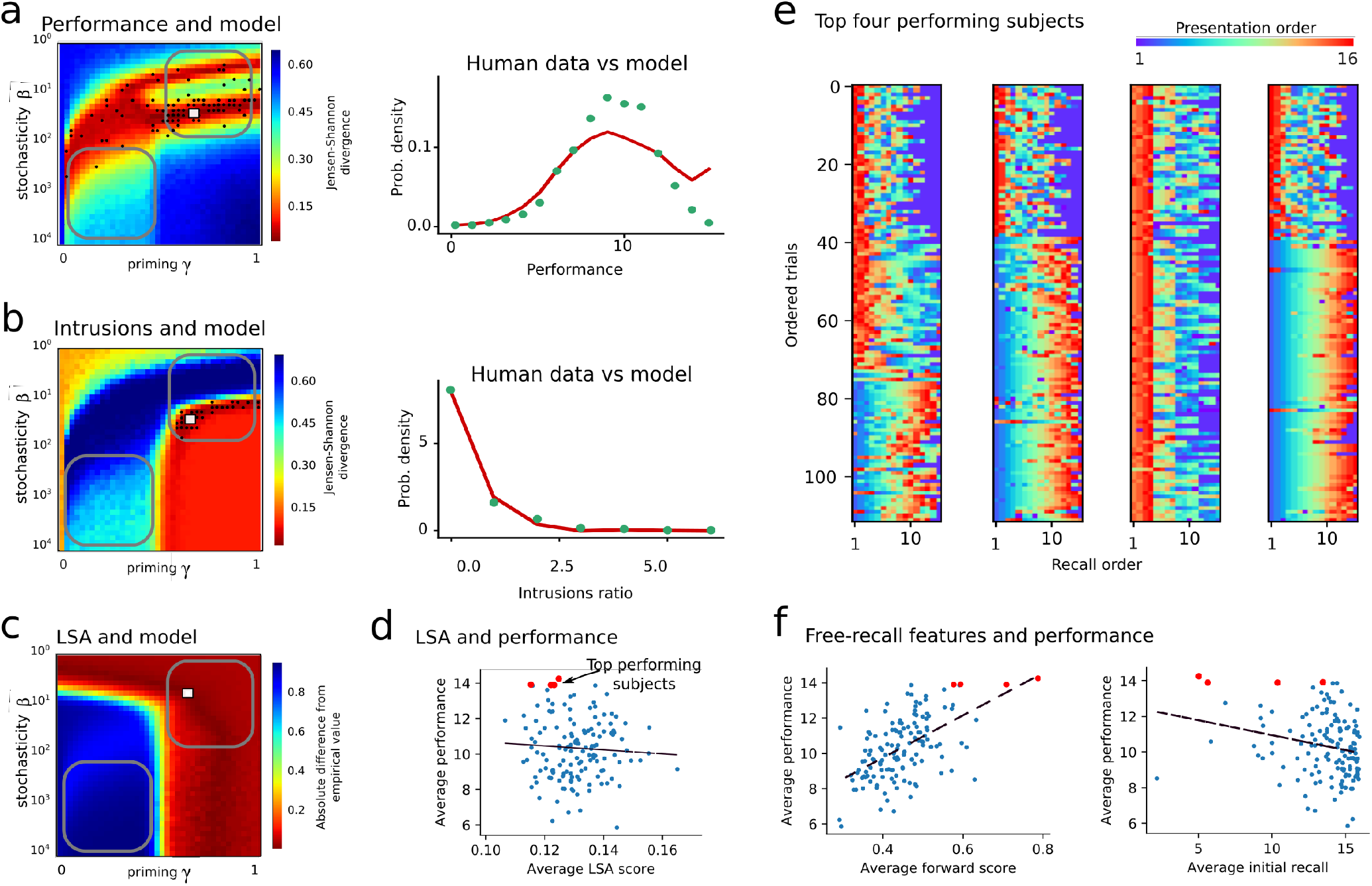
Analysis of free recall data. a) Comparison of human vs model free recall performance. Optimal performance of 16 is equal to the number of items presented. Left: JS-divergence between free-recall data and the model curves vs model parameters (*γ, β*). Lower values (red) indicate stronger agreement between data and model. White point corresponds to model distribution best approximating all free-recall data (performance distributions at these parameters shown to right). Black dots correspond to model distributions best approximating individual subjects recall data. Marked regions are those from Fig. 2h, approximately optimal for natural and free recall. b) As in A except using intrusion ratio instead of performance c) Comparison between the probability that transitions between retrieved items follow a high-LSA (Latent Semantic Analysis) similarity in the data, or a network connection in the model. Heatmap shows absolute difference between the probabilities for the model and the data. White dot indicates the parameters of minimal difference. d) Correlation between performance and average LSA score between retrieved items across all subjects (correlation coefficient *r* = −0.064, p-value= 0.45). e) Visualization of the recall statistics of top performing human subjects. Each plot represents shows 108 retrieval sessions (trials), ordered by trial number. In each trial (row) a variable number of items was retrieved (from 1 to 16). Retrieved items are shown in the order they were retrieved but colored by presentation order. f) Left: Performance vs forward score: the probability that a retrieval transition preserves the items presentation order (*r* = 0.47, p-value < 10^−11^). Right: Performance vs average presentation position of first item retrieved (*r* = 00.55, p-value= 0.0068).

We next asked whether variation in recall performance across subjects could be systematically explained. Although performance across subjects slightly correlated with variations in the model parameters fit to individual subjects (black dots in Figs. 3a to 3b), this did not appear to be a systematic explanation. Varying *γ* within the free-recall regime, which controls how strongly recall exploits the cognitive network, only weakly affected our model’s performance in free recall (Figs. 2e and 2h). Correspondingly, by approximating cognitive associations via high LSA scores, we found no significant correlation between mean subject performance and mean LSA scores between consecutively retrieved words (Fig. 3d). This suggests that top performance does not emerge simply via different priming and stochasticity parameters.

To gain insight into the factors driving free-recall performance we instead carefully inspected retrieval in the top performers. Curiously, the top performers retrieved the presented list following specific patterns Fig. 3f. Some subjects developed a retrieval pattern over the course of multiple trials while others jumped to it abruptly (31, 32). While it is known that better performing subjects tend to consecutively retrieve items that were initially presented nearby in time (the temporal contiguity effect (33)), the retrieval patterns of multiple top performers exhibited additional structure still. Top performers appeared to retrieve memory items in order, leading us to hypothesize that they were employing a different retrieval strategy from worse performers. In support of this hypothesis, the average number of forward transitions in free recall (the percentage of recall transitions that preserved the presentation order, Fig. 3f left) correlated with performance. Additionally, the true position of the first retrieved item anti-correlated with performance: better performers tended to initiate retrieval with earlier presented items (Fig. 3f right). Both of these features suggest ordering information may play an important role even in memory even in contexts where it is not required. This aligns with the overarching picture we have introduced that the cognitive states we visit during both free and natural memory tasks have an intrinsic ordering that our memory system might exploit for recall.

In the final section we show how our natural recall paradigm can also store and retrieve arbitrary serial order, upon considering more general and potentially realistic cognitive networks than we have so far explored.

### Encoding arbitrary sequences via auxiliary paths

The expressivity of path vectors is determined by how well sequences can be encoded as cognitive network paths. So far all cognitive states have corresponded to distinct items, but in general these may be a subset of the total states. If only some states represent items explicitly but these can be connected via paths through non-item auxiliary (aux) states, one could string items in flexible orders by connecting them via auxiliary paths (Fig. 4a). The item sequence is then retrieved by retracing this auxiliary path from its path vector (starting from a recall CUE state), reporting any item states passed along the way. To distinguish item from aux states in recall one can weight aux states by .5 when forming 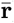 (Methods), so aux states will still be preferentially revisited in recall but distinguishable from item states: 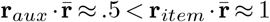. Thus, we let natural recall now refer to recall of paths in a more general cognitive network, allowing path vectors to recruit aux states to store arbitrary list orders, reflecting the top performers’ strategy in Fig. 3e.

**Fig.4.**
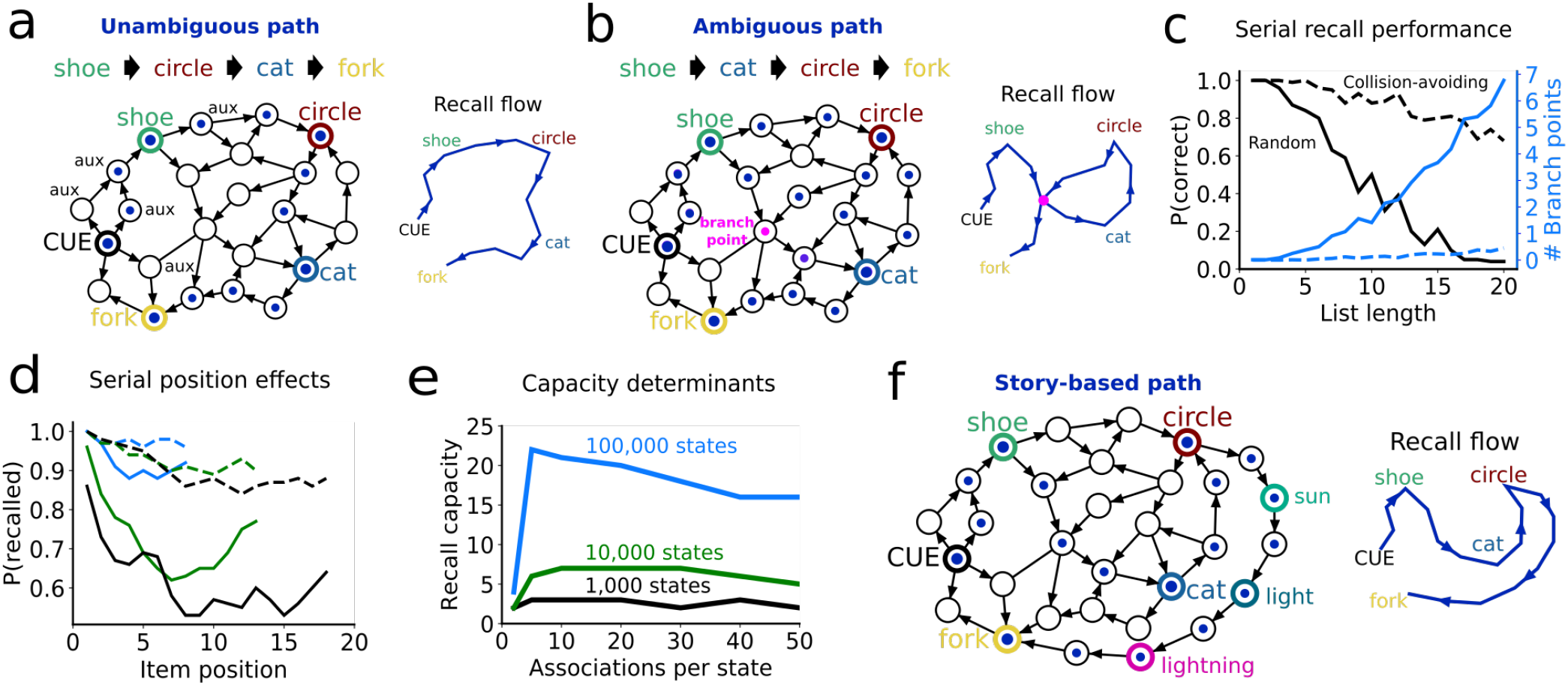
Sequence memory via auxiliary paths through general cognitive networks. a) Schematic generalized cognitive network with item (colored) and auxiliary non-item (black) states, target sequence to store, path encoding, and flow of recall dynamics (right) for unambiguous auxiliary path. Large dots correspond to weights of 1 in 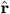, small dots .5. To recall the list, start at CUE and follow trail of inscribed dots along network, reporting items when large dots are reached. b) As in A, but for an ambiguous path containing one branch point (pink). c) Fraction of word lists recalled perfectly vs list length (black), using either random shortest vs collision-avoiding auxiliary paths, and the mean number of corresponding branch points (blue). 100 trials were performed on a network with 10,000 total states with 100 item states, averaging 10 associations per state; trials terminated in the absence of a subsequent primed association or when twice as many words were recalled as had been presented (counting repeats). d) Probability a given item was recalled vs its position in the presentation list, for random shortest (solid) and collision-avoiding (dashed) algorithms for constructing auxiliary paths, different colors indicate different list lengths. e) Recall capacity (longest list with a 50% chance of perfect retrieval) for different network sizes and mean associations per state (100 item states in all cases). f) As in B except using auxiliary path connecting CIRCLE to FORK via a “storyline” following semantically related item states through a separate section of the network (not shown), with no branch points.

Even if (*β, γ*) are tuned for natural recall, serial recall errors arise if the auxiliary path cannot be unambiguously recovered from its states. If any state primed by (included in) the path vector has more than one primed association, this state becomes a “branch point” (Fig. 4b), as the next state recalled may differ across trials. When the branch point in the path representation of [SHOE, CAT, CIRCLE, FORK] is reached in Fig. 4b, the state toward FORK will follow as often as the state toward CAT, so CAT and CIRCLE can be skipped in recall (Fig. 4b). Branch points emerge when the auxiliary path collides with itself, yielding spurious connections that can bypass correct items during recall. Which items are skipped and how often depends on network structure and how the auxiliary path is built.

Even in a random network, branch points in auxiliary paths yield errors resembling human recall. Using a large network with random sparse associations, one CUE state and 100 random states assigned to items, we tested storing random item lists via auxiliary paths. During list presentation we formed an auxiliary path by linking each consecutive item pair (starting at CUE) via a random shortest path between the item states, i.e. the cognitive state is steered along the network as directly as possible to the next item state. To focus on branch point errors we considered the natural recall regime and *N* → ∞. Branch points increased and serial recall worsened with list length due to an increased chance of collisions in longer auxiliary paths (Fig. 4c), resembling the limited recall capacity of humans (34). Early and later items appeared more than middle items in recall, reflecting primacy and recency (Fig. 4d) (28, 35): branch point opportunities increase as one moves from early to middle items, causing primacy; and whereas one can get stuck downstream of middle items, this is less likely for later items, which are thus skipped less, yielding a recency effect similar to experimental observations (although other mechanisms may influence recency as well). Serial recall capacity increased with network size and with sparse but nonzero state associations (Fig. 4e); too many associations increase collisions in auxiliary paths; too few limit paths for linking item states. Thus, even with a random network recall reflects human errors and is quantitatively shaped by network structure.

Crucially, recall depends on how auxiliary paths are built. When we instead built auxiliary paths in the same networks so as to explicitly avoid collisions, branch points decreased and recall capacity increased dramatically (Figs. 4c to 4d). In principle auxiliary paths could also treat extra item states themselves as aux states, weighting them by .5 in 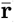). E.g. one could build a bridge from CIRCLE to FORK via CIRCLE → SUN → LIGHT → LIGHTNING → FORK (Fig. 4f), to explicitly decrease the chance of colliding with CAT. This resembles how humans can improve recall by using auxiliary storylines to connect presented items (24). If naturally associated items like SUN and LIGHT are closer on the network than unrelated items, building paths that recruit these could yield fewer branch points, suggesting a specific mechanism for how auxiliary storylines improve recall.

## Discussion

We introduced a two-level model for storing flexible sequential memories. An event or item sequence is mapped to a path through a network of cognitive states, whereupon the states’ neural representations are summed into a *path vector* from which the sequence can later be retrieved. Networks of hidden states have found success in accounting for animal behavior and neural activity (36, 37) and human free recall (6, 38, 39); however, it is unknown how sequences of states are stored (e.g. to repeat back a series of stimuli or actions), possibly retained in a unique compact representation, and retrieved. Unlike previous models, we allow for auxiliary states that do not directly correspond to external observables, which can be flexibly recruited to encode novel serial relationships. Our model demonstrates how state sequences that are more closely aligned with current network pathways, referred to as “natural” sequences, would be stored more effectively since fewer auxiliary states are required. We also explain how this helps with subsequent retrieval.

Since path vectors are created by simple summation, each neuron need only remember the sum total of its own activity over the sequence presentation. This represents a bio-plausible demand, requiring neither additional timestamping operations, persistent dynamics, nor Hebbian synaptic changes, and which might instead be implemented cellularly (40–42). Our model thus is fundamentally distinct from existing models of distributed sequence memory, such as the temporal context model (8) or state-of-the-art vector symbolic architectures (9), which encode serial order by timestamping item representations via multiplicative or convolutional operations, which may require more complex biological mechanisms. Hippocampus, known for fast plasticity and sequence coding (43, 44), could be well suited to rapidly construct path vectors. Neocortex, thought to learn knowledge structures and associations (45), would be a natural substrate for the cognitive state association network. Storage and readout of episodes could potentially unfold via hippocampal-cortical interactions.

Path vectors exhibit several advantageous coding properties. First, information about items and their order is maximally distributed (all neurons are equal), making the code robust to loss of arbitrary neuron groups. Second, path vectors have fixed dimension for all sequence lengths so no preallocation of memory space is required, allowing them to encode ongoing streams of experience with no pre-known endpoint; instead, encoding degrades gracefully with episode length. Third, arbitrary sequences do not have unique representations but admit encoding by flexible auxiliary paths, allowing them to reroute around lost or unavailable cognitive states. Connecting items with paths has been demonstrated to be effective for encoding the serial order of spatial locations, while path crossings in 2-D space were found to impair recall. ((46)). Our work transposes this idea from 2-D space to network topologies linking general cognitive states. As a result, our model predicts that memory would be hindered when the same or comparable cognitive state is visited again while an item list is presented, even if the list contains no repetitions. This prediction may be confirmed using neuroimaging.

Although random networks reproduced certain human recall patterns (Figs. 3 and 4), networks of associations among real cognitive states likely have more structure. States representing locations, for instance, may be connected via lattice-like associations (42, 46, 47); and more general state associations may be shaped by individual experience. Network topology in turn constrains which paths can be built by which algorithms: size and sparsity of random networks affect the length of non-branching paths realizable by linking random shortest paths between item states (Fig. 4e), and in fixed networks path-building algorithms suggestive of story-like mnemonics increase capacity (Figs. 4c and 4f). How networks representing associations among more realistic story elements would support bioplausible path-building is an open question, which our work sets the stage to quantitatively address.

For simplicity we modeled cognitive states with nearly orthogonal vector representations. Deviations from orthogonality, however, are implicated in generalization and abstraction, two key aspects of cognition (48, 49). Two cognitive states might have a higher neural dot product if they encode related over dissimilar items, for example. In the context of learning this could aid in generalization but also introduces the potential for confusing them, a fundamental trade-off surrounding shared representations in parallel information processing (50). We anticipate that our results will hold for deviations from orthogonality as long as they are not so severe that states become confusable when summed into the path vector. For example, it is known that summing non-orthogonal word embeddings produces a suitable representation of sentences for many tasks (51), implying that simple vector sums can capture both implicit ordering information in training data as well as semantic associations. A fascinating future direction will be to investigate how the topology of cognitive state networks and the geometry of state representations might be combined to serve tasks that require both memory and abstraction.

## Methods

### Neural representations of cognitive states

For all simulations and analyses, vector representations {**r**_*i*_} of cognitive states {*v*_*i*_} are independent *N* -dimensional random vectors with i.i.d. components sampled from a zero-mean Gaussian with variance 1*/N*.

### Free recall dataset

The data presented in this article were gathered in M. Kahana’s lab as part of the Penn Electrophysiology of Encoding and Retrieval Study (details of the experiments may be found in (29)). Participants gave their permission in accordance with the University of Pennsylvania’s IRB procedure and were paid for their time. We examined the data from the 141 individuals (aged 17 to 30) who completed the first phase of the experiment, which consisted of 7 experimental sessions. Each session lasted about 1.5 hours and consisted of 16 lists of 16 words given one at a time on a computer screen. Each study list was immediately followed by a free recall testing. Words were chosen from a pool of 1638. There was a 1500 ms wait before the first word showed on the screen for each list. Each item was shown on the screen for 3000 milliseconds, followed by jittered 800 - 1200 millisecond inter-stimulus intervals (uniform distribution). Following the final item on the list, there was a 1200 - 1400 ms jittered delay, followed by 75 seconds for participants to try to remember any of the previously presented items.

### Data analysis

We compared our model with free recall data via:

#### Performance

The number of unique correct (shown during the presentation phase) items reported during recall.

#### Intrusion ratio

During the retrieval phase subjects often reported items that were not presented, called *intrusions*. The intrusion ratio is the number of intrusions before the last retrieved item divided by the performance (number of correct words retrieved).

#### LSA metric

We considered the Latent Semantic Analysis (LSA) score between any two words of the 1638 words that were presented to subjects during the experiment. For each word we considered it semantically related to another if the LSA score between any the first and the second was in the top 100(1 − *p*) percentile of the probability distribution of LSA scores for that word. Here *p* matches the sparsity of the model simulations *p* = 0.01. We then computed the probability that a transition, during retrieval, followed a semantic link. We performed the same analysis on the model, where we computed the probability that a transition between two visited words by the retrieval dynamics would follow a link in the cognitive network.

### Model simulation and analysis

Each simulation of the model correspond to a trial of an experimental paradigms, comprising a presentation and a recall phase (Fig. 2a).

#### Free recall

In free recall first a set of M=1000 random vectors in N=500 dimensions is constructed. Each vector 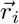 corresponds to one of M items (*i* ∈ 1..*M*). Components are i.i.d. Gaussian with zero mean and variance 1/500. Next, a random binary directed graph adjacency matrix *W* (Erdos-Renyi) of size *M × M* with sparsity 0.01 (each item is connected to any other with probability p=0.01) is generated. This graph *W* defines which memory items are connected in what refer to as network of cognitive associations. These objects are fixed across trials. In the presentation phase of each simulation *L* = 16 items are randomly chosen and presented in a random order, we term the set of these items *k*_1_, …, *k*_*L*_. Presentation leads to the formation of a path vector 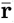 (Fig. 2a) i.e. the sum of the presented states’ representations.

The presentation phase followed a recall phase. In free recall this was initialized at a random correct item, then followed Equation (2). The parameters *γ, β* capture respectively the amount of noise and the strength of the influence of the path-vector on the recall process. The recall phase progresses for *T* = 100 retrieval steps, with items originally included in the presentation phase counted as correct. If all correct items were retrieved within T steps, we let *t*_*final*_ indicate the time the final correct item was retrieved, marking the effective end of the recall phase; all items visited at subsequent steps were discarded. This is to resemble the experimental paradigm where subjects were given 75 seconds to retrieve items, but where often all items were recalled before 75 seconds and the trial terminated. Before the end of retrieval some items which were not presented were often visited. The number of unique items visited in such way was deemed *N*_*intrusions*_. The metrics displayed in Figs. 2d to 2g can be directly defined in terms of the above quantities:

- Recall performance: this is *N*_*retrieved*_;
- Retrieval speed: this is given by *N*_*retrieved*_*/t*_*final*_;
- Intrusions rate: this is given by *N*_*intrusions*_*/t*_*final*_.
- Ranked similarity between items (LSA comparison): for consecutively retrieved correct items, this is given by the the ranked probability of transition between the current item and the next, among all probabilities of transitions 𝒫(*j* → *i*) from the current item *k*^*j*^ and any other presented one.

We reported these quantities as averaged across multiple simulations *N*_*s*_ = 1000 for each parameter pair (*β, γ*) in Figs. 2d to 2g and Figs. 3a to 3c.

#### Natural recall

Identical to free recall expect that:

- Presented items where not drawn purely at random. Instead, for each simulation a first item *k*_*ini*_ was randomly chosen and then the graph *W* of associations between items/states was followed choosing a random link from the initial item to a subsequent one and so on. The string of visited *L* = 16 items was then used as presented items.
- In the recall phase the initial item was initialized to be the correct first item *k*_*ini*_. Recall followed the same rules as above except items had to be retrieved in the same presentation order to count as correct. Omissions were allowed, so for example if items *k*^1^…*k*^1^6 were presented a possible recall was *k*_1_, *k*_2_, *k*_4_, intrusion, *k*_9_, intrusion, *k*_12_.
- The number of words retrieved was similarly computed, and *t*_*final*_ marked the first occurrence of the last retrieved items, as in free recall.

### Path vectors with auxiliary states

To form a path vector 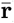 that includes a set of item states *V*_*item*_ and a set of auxiliary states *V*_*aux*_ we weight the item states by 1 and the aux states by .5:

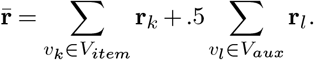

### Arbitrary sequences and recall strategies

Here (Fig. 4) we ran all simulations in the large-*N* limit in the natural recall regime. I.e. we assumed perfect ability to read out whether a state was contained in the path vector, such that the only recall errors were those caused by reaching branch points, which leads to ambiguity over which state to visit next. When there were *Z* downstream states included in the path vector, the chance of visiting any of them was 1*/Z*. Identifying a primed downstream state in this limiting case amounts to checking each possible downstream states’ dot product with **r**, which requires 𝒪(*N* ⟨*Q*⟩ *L*^*^) operations, where ⟨*Q*⟩ is the average out-degree of the network and *L*^*^ the number of retrieval steps.

In the “random” strategy, an auxiliary path encoding a sequence of items *k*_1_, …, *k*_*L*_ was built as follows. All shortest directed paths linking pairs consecutive items were found, including a link between the CUE state and 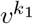. For each consecutive item pair one shortest path linking that item pair was chosen at random, and all states included in the chosen shortest paths were summed into the path vector.

In the “collision-avoiding” strategy, for each consecutive item pair (including CUE), all directed paths up to a max length of 15 states were identified, and a preliminary complete auxiliary path was constructed by randomly selecting one directed path linking each item pair. If the auxiliary path contained branch points, it was subsequently “untangled” by iteratively (1) identifying all branch points, (2) for each branch point identifying the item pairs whose connecting paths collided to create the branch point, (3) randomly resampling a new path for each of these item pairs. This procedure was repeated until there were either no more branch points or there were 25 failed attempts to decrement the branch points in the auxiliary path. This procedure demonstrates proof-of-concept differences between random vs collision-avoiding paths on recall performance, but we do not hypothesize that this algorithm is used by the brain.

## Acknowledgments

This research was supported by the espresso machine in the University of Washington Computational Neuroscience Center. We would like to thank Alison Weber, Matthew Farrell, Kamesh Krishnamurthy, Tankut Can, Julia Steinberg, Lee Susman, Misha Katkov, Misha Tsodyks, and Francesco Fumarola for fruitful discussions in the development of this manuscript.

## References

1. John E Lisman and Marco A Idiart. Storage of 7+/-2 short-term memories in oscillatory subcycles. Science, 267(5203):1512–1515, 1995.

2. Margaret F. Carr, Shantanu P. Jadhav, and Loren M. Frank. Hippocampal replay in the awake state: a potential substrate for memory consolidation and retrieval. Nature Neuroscience, 14(2):147–153, February 2011. ISSN 1097-6256. doi: 10.1038/nn.2732.

3. Jon W Rueckemann, Marielena Sosa, Lisa M Giocomo, and Elizabeth A Buffalo. The grid code for ordered experience. Nature Reviews Neuroscience, 22(10):637–649, 2021.

4. Rodrigo Laje and Dean V Buonomano. Robust timing and motor patterns by taming chaos in recurrent neural networks. Nature neuroscience, 16(7):925–933, 2013.

5. Kanaka Rajan, Christopher D Harvey, and David W Tank. Recurrent network models of sequence generation and memory. Neuron, 90(1):128–142, 2016.

6. Vezha Boboeva, Alberto Pezzotta, and Claudia Clopath. Free recall scaling laws and short-term memory effects in a latching attractor network. Proceedings of the National Academy of Sciences, 118(49), 2021.

7. Michael E Hasselmo and Bradley P Wyble. Free recall and recognition in a network model of the hippocampus: simulating effects of scopolamine on human memory function. Behavioural Brain Research, 89(1–2):1–34, December 1997. ISSN 0166-4328. doi: 10.1016/S0166-4328(97)00048-X.

8. Marc W Howard and Michael J Kahana. A distributed representation of temporal context. Journal of mathematical psychology, 46(3):269–299, 2002.

9. Pentti Kanerva. Hyperdimensional computing: An introduction to computing in distributed representation with high-dimensional random vectors. Cognitive computation, 1(2):139– 159, 2009. Publisher: Springer.

10. Matthew M Botvinick and David C Plaut. Short-term memory for serial order: a recurrent neural network model. Psychological review, 113(2):201, 2006.

11. Zachary C Lipton. The mythos of model interpretability: In machine learning, the concept of interpretability is both important and slippery. Queue, 16(3):31–57, 2018.

12. Gary Gillund and Richard M Shiffrin. A retrieval model for both recognition and recall. Psychological review, 91(1):1, 1984.

13. Richard Nevill Astley Henson. Short-term memory for serial order. PhD thesis, University of Cambridge UK, 1996.

14. Donald Laming. Failure to recall. Psychological Review, 116(1):157, 2009.

15. Melissa Lehman and Kenneth J Malmberg. A buffer model of memory encoding and temporal correlations in retrieval. Psychological Review, 120(1):155, 2013.

16. Sandro Romani, Itai Pinkoviezky, Alon Rubin, and Misha Tsodyks. Scaling laws of associative memory retrieval. Neural computation, 25(10):2523–2544, 2013.

17. Sen Song and Larry F Abbott. Cortical development and remapping through spike timing-dependent plasticity. Neuron, 32(2):339–350, 2001.

18. Stefan Klampfl and Wolfgang Maass. Emergence of dynamic memory traces in cortical microcircuit models through stdp. Journal of Neuroscience, 33(28):11515–11529, 2013.

19. Randal A Koene and Michael E Hasselmo. First-in–first-out item replacement in a model of short-term memory based on persistent spiking. Cerebral Cortex, 17(8):1766–1781, 2007.

20. Joel Walters and Yuval Wolf. Language proficiency, text content and order effects in narrative recall. Language Learning, 36(1):47–64, 1986. Publisher: Wiley Online Library.

21. Joshua J. Diehl, Loisa Bennetto, and Edna Carter Young. Story recall and narrative coherence of high-functioning children with autism spectrum disorders. Journal of abnormal child psychology, 34(1):83–98, 2006. Publisher: Springer.

22. Monika Fludernik. Towards a’natural’narratology. jRoutledge, 2002.

23. Peter Hühn, Jan Christoph Meister, John Pier, and Wolf Schmid. Handbook of Narratology. Walter de Gruyter GmbH & Co KG, October 2014. ISBN 978-3-11-031646-9. Google-Books-ID: v9fmBQAAQBAJ.

24. Gordon H Bower and Michal C Clark. Narrative stories as mediators for serial learning. Psychonomic Science, 14(4):181–182, 1969.

25. Pietro Berkes, Gergo? Orbán, Máté Lengyel, and József Fiser. Spontaneous cortical activity reveals hallmarks of an optimal internal model of the environment. Science, 331(6013): 83–87, 2011.

26. Matthew A Wilson and Bruce L McNaughton. Dynamics of the hippocampal ensemble code for space. Science, 261(5124):1055–1058, 1993.

27. Sanjoy Dasgupta, Charles F Stevens, and Saket Navlakha. A neural algorithm for a fundamental computing problem. Science, 358(6364):793–796, 2017.

28. Michael Jacob Kahana. Foundations of human memory. OUP USA, 2012.

29. Jonathan F Miller, Christoph T Weidemann, and Michael J Kahana. Recall termination in free recall. Memory & cognition, 40(4):540–550, 2012.

30. Weston A Bousfield and Charles Hill W Sedgewick. An analysis of sequences of restricted associative responses. The Journal of General Psychology, 30(2):149–165, 1944.

31. Sandro Romani, Mikhail Katkov, and Misha Tsodyks. Practice makes perfect in memory recall. Learning & Memory, 23(4):169–173, 2016.

32. Michelangelo Naim, Mikhail Katkov, Stefano Recanatesi, and Misha Tsodyks. Emergence of hierarchical organization in memory for random material. Scientific reports, 9(1):1–10, 2019.

33. Per B Sederberg, Jonathan F Miller, Marc W Howard, and Michael J Kahana. The temporal contiguity effect predicts episodic memory performance. Memory & cognition, 38(6):689– 699, September 2010. ISSN 1532-5946. doi: 10.3758/MC.38.6.689.

34. George A Miller. The magical number seven, plus or minus two: Some limits on our capacity for processing information. Psychological review, 63(2):81, 1956.

35. Eddy J Davelaar, Yonatan Goshen-Gottstein, Amir Ashkenazi, Henk J Haarmann, and Marius Usher. The demise of short-term memory revisited: empirical and computational investigations of recency effects. Psychological review, 112(1):3, 2005.

36. Adam J Calhoun, Jonathan W Pillow, and Mala Murthy. Unsupervised identification of the internal states that shape natural behavior. Nature neuroscience, 22(12):2040–2049, 2019.

37. Stefano Recanatesi, Ulises Pereira-Obilinovic, Masayoshi Murakami, Zachary Mainen, and Luca Mazzucato. Metastable attractors explain the variable timing of stable behavioral action sequences. Neuron, 2021.

38. Francesco Fumarola. A diffusive-particle theory of free recall. Advances in cognitive psychology, 13(3):201, 2017.

39. Michelangelo Naim, Mikhail Katkov, Sandro Romani, and Misha Tsodyks. Fundamental law of memory recall. arXiv preprint 1905.02403, 2019.

40. Alexei V Egorov, Bassam N Hamam, Erik Fransén, Michael E Hasselmo, and Angel A Alonso. Graded persistent activity in entorhinal cortex neurons. Nature, 420(6912):173– 178, 2002.

41. Wei Zhang and David J Linden. The other side of the engram: experience-driven changes in neuronal intrinsic excitability. Nature Reviews Neuroscience, 4(11):885–900, 2003.

42. Rich Pang and Adrienne L Fairhall. Fast and flexible sequence induction in spiking neural networks via rapid excitability changes. Elife, 8:e44324, 2019.

43. David J Foster and James J Knierim. Sequence learning and the role of the hippocampus in rodent navigation. Current opinion in neurobiology, 22(2):294–300, 2012.

44. Katie C Bittner, Aaron D Milstein, Christine Grienberger, Sandro Romani, and Jeffrey C Magee. Behavioral time scale synaptic plasticity underlies ca1 place fields. Science, 357 (6355):1033–1036, 2017.

45. Patricia S Goldman-Rakic. Topography of cognition: parallel distributed networks in primate association cortex. Annual review of neuroscience, 11(1):137–156, 1988.

46. Fabrice BR Parmentier, Greg Elford, and Murray Maybery. Transitional information in spatial serial memory: Path characteristics affect recall performance. Journal of Experimental Psychology: Learning, Memory, and Cognition, 31(3):412, 2005.

47. Margaret F Carr, Shantanu P Jadhav, and Loren M Frank. Hippocampal replay in the awake state: a potential substrate for memory consolidation and retrieval. Nature neuroscience, 14(2):147–153, 2011.

48. Silvia Bernardi, Marcus K Benna, Mattia Rigotti, Jérôme Munuera, Stefano Fusi, and C Daniel Salzman. The geometry of abstraction in the hippocampus and prefrontal cortex. Cell, 183(4):954–967, 2020.

49. SueYeon Chung and LF Abbott. Neural population geometry: An approach for understanding biological and artificial neural networks. arXiv preprint 2104.07059, 2021.

50. Giovanni Petri, Sebastian Musslick, Biswadip Dey, Kayhan Özcimder, David Turner, Nesreen K Ahmed, Theodore L Willke, and Jonathan D Cohen. Topological limits to the parallel processing capability of network architectures. Nature Physics, 17(5):646–651, 2021.

51. Sanjeev Arora, Yingyu Liang, and Tengyu Ma. A simple but tough-to-beat baseline for sentence embeddings. In International conference on learning representations, 2017.

